# Role of mast cells in endomicrobial sepsis revisited: Mast cell-deficient mice show normal immunological protection

**DOI:** 10.1101/2025.03.07.642033

**Authors:** Thorsten B. Feyerabend, Fabienne Schochter, Alpaslan Tasdogan, Maren Bechberger, Sébastien Boutin, Hans-Reimer Rodewald

## Abstract

*Kit*-mutant mice are highly susceptible to polymicrobial sepsis elicited by cecal ligation and puncture (CLP). This vulnerability has been attributed to the mast cell deficiency of *Kit* mutants, suggesting key roles of mast cells in antibacterial defense. We show that mice lacking mast cells but wild-type for *Kit* are as resistant to sepsis as mast cell-proficient mice, excluding mast cells as protective factor. Induction of sepsis by direct injection of intestinal microbiota instead of surgical gut perforation revealed comparable protective responses of *Kit*-deficient and *Kit* wild-type mice, indicating intact antibacterial immunity in the absence of Kit. Notably, cecal contents of *Kit*-mutant mice contained 1000-fold greater *Escherichia coli* colony-forming units compared to wild-type mice. Consistently, 16S rRNA gene sequencing revealed a broad compositional shift characterized by enrichment of Enterobacteriaceae and other taxa associated with inflammatory intestinal states, in keeping with intestinal dysmotility in *Kit*-mutant mice. Thus, upon intestinal puncture, overrepresentation of pathogenic bacteria led to a more severe infection of *Kit* mutants compared to wild-type controls. Hence, the susceptibility of *Kit*-mutant mice to sepsis is caused by enteral dysbiosis, not mast cell deficiency. These findings highlight the importance of considering genotype-dependent effects on endogenous microbiota composition in CLP models. Collectively, our results show that immunological resistance to sepsis is independent of mast cells.

## Introduction

30 years ago, two hallmark papers studying acute peritoneal infections reported evidence for protective functions of mast cells in anti-bacterial defense^1,2^. These experiments were performed in mice carrying inactivating mutations in the gene encoding the receptor tyrosine kinase Kit (*Kit^W/Wv^* mice) which, at the time, were the widely used model of mast cell deficiency owing to the dependency of mast cell development on Kit expression^3^. Both reports showed that *Kit^W/Wv^* mice were highly susceptible to peritoneal infections compared with *Kit* wild-type mice. Based on mast cell reconstitution and investigation of associated cytokines, the central conclusion was that the absence of mast cells in *Kit^W/Wv^* mice, and in particular the absence of mast cell-derived tumor necrosis factor alfa (TNFα), was responsible for the observed immunodeficiency^1,2^. These papers were very influential, and mast cells have since been viewed as key effectors in bacterial defense^4^.

At the time, it was well established that *Kit* mutations affect many cell lineages and tissues in developing and adult mice beyond the defect in mast cells, and hence it remained ambiguous whether or not a phenotype observed in *Kit*-mutant mice was indeed due to the absence of mast cells. With the advent of *Kit* mutation-independent mouse models of mast cell deficiency^5–8^, which are by comparison to *Kit* mutants highly specific experimental models^9–13^, mast cell functions can be probed more conclusively in vivo. It has since become possible to independently confirm, or disprove, what had been concluded earlier on mast cell physiology based on work in *Kit* mutants. Around 15 years into this process of re-evaluation of mast cell functions, a long list of suggested functions have not been reproduced when revisited in *Kit*-independent models (for discussions and references, see^9–13^).

Here, we have re-addressed the important question whether mast cells are involved in immunity to acute peritonitis and sepsis. We confirmed the susceptibility of *Kit*-mutant mice (*Kit^W/Wv^*) to sepsis^1^ in the cecal ligation and puncture (CLP) model^14^. However, comparison of *Kit* wild-type mice with (*Cpa3^+/+^*) or without (*Cpa3^Cre/+^*) mast cells^6^ showed that the outcome of such infections was independent of mast cells. Furthermore, *Kit^W/Wv^* mice exhibited the same susceptibility as *Kit^+/+^* mice upon direct injection of intestinal bacteria, pointing at a defect in *Kit^W/Wv^* mice related to the intestinal injury, or the release of endogenous flora from the cecum in the CLP model. Finally, we identified markedly higher *Escherichia coli* (*E. coli*) colony-forming units in the cecum of *Kit^W/Wv^* mice compared to *Kit^+/+^* mice. At greater resolution, 16S rRNA gene sequencing revealed a pronounced dysbiosis characterized by enrichment of inflammation-associated bacterial taxa in *Kit^W/Wv^* mice compared to wild-type controls. This dysbiosis is unrelated to mast cell deficiency because mice with (*Cpa3^+/+^*) or without (*Cpa3^Cre/+^*) mast cells had comparable microbiota based on 16S rRNA sequencing. The altered microbial composition in *Kit^W/Wv^* mice, explains their more severe infection challenge during CLP. Consistent with a microbiota-driven effect, co-housing neutralized the susceptibility of *Kit*-mutant mice to CLP-induced sepsis. Our data suggest that the dysbiosis of *Kit^W/Wv^* mutants has been misinterpreted as an immunodeficiency. Collectively, we provide evidence that mast cells are dispensable for protection against polymicrobial sepsis and that susceptibility of *Kit*-mutant mice to CLP is driven by elevated microbiota pathogenicity.

## Results

### Mast cells are dispensable for survival of cecal ligation and puncture-induced polymicrobial sepsis

We subjected mast cell-deficient *Cpa3^Cre/+^* mice^6^ on the C57BL/6 background (*Cpa3^Cre/+^*), and their wild-type littermates (*Cpa3^+/+^*) to the surgical model of cecal ligation and puncture^14^. Under mild sepsis conditions, induced by a 50% ligation of the cecum (tying at the middle between the base and the pole of the cecum) and a single puncture of the cecum with a 25-gauge (25 G) needle, all wild-type mice (*Cpa3^+/+^*) survived (Fig. 1A). Interestingly, all mast cell-deficient *Cpa3^Cre/+^* mice also survived (Fig. 1A). To explore whether a role for mast cell would become evident under more severe conditions we induced CLP using a 22 G needle (Fig. 1B). The survival (Methods) within two weeks after surgical puncture dropped to about half of the animals for both the *Cpa3^+/+^* and *Cpa3^Cre/+^* mice (Fig. 1B). We analyzed large cohorts of about n = 50 for each genotype, and found no difference between both groups (P = 0.23), indicating that the mortality in cecal ligation and puncture sepsis was independent of mast cells. These results were surprising given that previous publications using *Kit*-mutant, mast cell-deficient mice concluded that there was a fundamental requirement for mast cells in sepsis survival^1,2^. As a control, we therefore included *Kit^W/Wv^* mice in our studies. Even under mild sepsis conditions, which all *Kit* wild-type (*Cpa3^+/+^* or *Cpa3^Cre/+^*) mice survived, 75% of *Kit^W/Wv^* mice died (Fig. 1A), and in severe sepsis all *Kit^W/Wv^* mice succumbed the treatment within two days (Fig. 1B). *Kit^W/Wv^* mice are F1 offspring from WB-*Kit^W/+^* and B6-*Kit^Wv/+^* parents, hence they have a genetic F1 hybrid (WBB6F1) background^15^ (we refer here to WBB6F1*-Kit^W/Wv^* mice as *Kit^W/Wv^* mice). To exclude genetic background differences as a cause of the higher susceptibility of *Kit^W/Wv^* mice compared to *Cpa3^Cre/+^* mice, we repeated the CLP experiments using all mice on the WBB6F1 background (WBB6F1-*Cpa3^+/+^,* WBB6F1-*Cpa3^Cre/+^* and *Kit^W/Wv^*). On this background, mast cell-proficient WBB6F1-*Cpa3^+/+^* and mast cell-deficient WBB6F1-*Cpa3^Cre/+^* mice were both even more resistant to CLP than on the B6 background (Fig. 1C), which is consistent with an earlier report^16^, and hybrid vigor. In total, 83% of WBB6F1-*Cpa3^+/+^* and 74% of WBB6F1-*Cpa3^Cre/+^* mice survived (P = 0.24), while again all *Kit^W/Wv^* mice succumbed to sepsis. These experiments demonstrate that mast cells are not involved in sepsis resistance, yet *Kit* mutants are highly susceptible.

**Figure 1.**
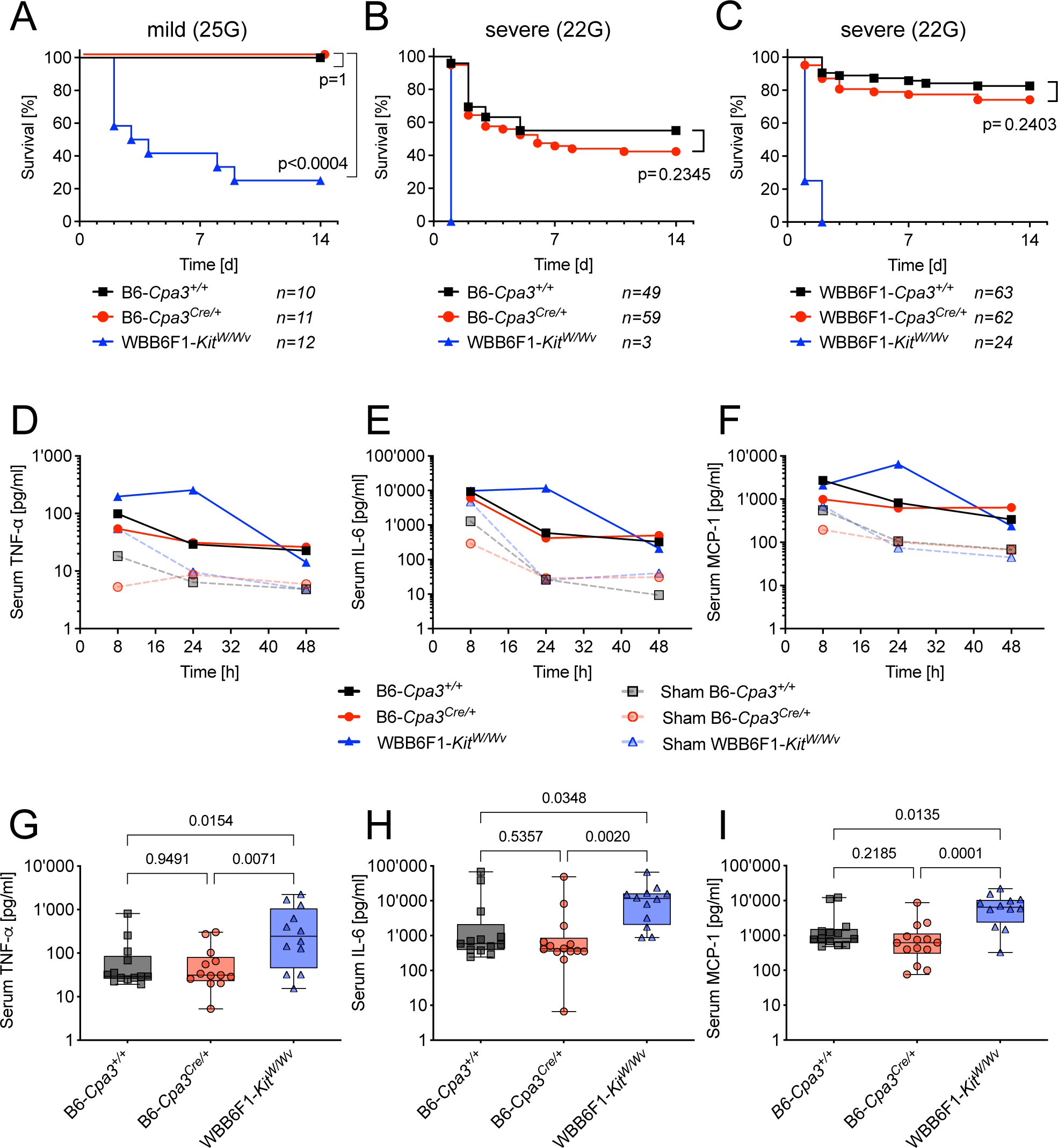
*Kit^W/Wv^* mutant mice, but not mast cell-deficient *Cpa3^Cre/+^* mice, are susceptible for cecal ligation and puncture sepsis. (A). Survival of B6-*Cpa3^+/+^* (100% of n = 10), B6-*Cpa3^Cre/+^* (100% of n = 11), and WBB6F1-*Kit^W/Wv^* (25% of n = 12) mice to mild (25 G needle) cecal ligation and puncture. WBB6F1-*Kit^W/Wv^* mice vs. B6-*Cpa3^+/+^* (P < 0.0004). (B). Survival of B6-*Cpa3^+/+^* (55% of n = 49), B6-*Cpa3^Cre/+^* (42% of n = 59), and WBB6F1-*Kit^W/Wv^* (0% of n = 3) mice under severe (22 G needle) cecal ligation and puncture. B6-*Cpa3^+/+^* vs. B6-*Cpa3^Cre/+^* P = 0.2345. WBB6F1-*Kit^W/Wv^* vs. B6-*Cpa3^+/+^* P < 0.0001. (C). Survival of WBB6F1-*Cpa3^+/+^*(82.5%, n = 63), WBB6F1-*Cpa3^Cre/+^* (74.2%, n = 62), and WBB6F1-*Kit^W/Wv^* (0%, n = 24) mice to severe (22 G needle) cecal ligation and puncture. P values for curve comparisons in A-C were calculated using the Mantel-Cox Log-rank test. (D-F). Kinetics (in hours) of median serum concentrations for TNFα (D), IL-6 (E), and MCP-1 (F) in B6-*Cpa3^+/+^*, B6-*Cpa3^Cre/+^*, and WBB6F1-*Kit^W/Wv^* mice after severe (22-G needle) cecal ligation and puncture (solid lines), or after sham operations (same procedure but without cecal ligation and puncture) (dashed lines). (G-I). From the kinetic shown in D-F, box plots were drawn for the 24-h time point indicating serum concentrations for TNFα, IL-6, and MCP-1 in B6-*Cpa3^+/+^*, B6-*Cpa3^Cre/+^* and WBB6F1-*Kit^W/Wv^* mice. Each dot represents an individual mouse. The boxes extend from the 25th to 75th percentiles and the median is indicated. Whiskers range from minimum to maximum. P values were calculated on log-transformed values by one-way ANOVA with Tukey’s correction for multiple comparisons.

Because absence of mast cells in *Kit^W/Wv^* mice has been implicated in impaired cytokine responses in sepsis, notably in TNFα^1,2^, we next analyzed the inflammatory cytokine response. Serum levels of TNFα, IL-6, and MCP-1 were measured at three different time points after CLP (Fig. 1D-F). Mast cell-proficient *Cpa3^+/+^* and mast cell-deficient *Cpa3^Cre/+^* mice showed very similar time course and amounts of cytokine production. The serum levels for each of the three cytokines was maximal after 8 h, and declined until 24 h without any significant further change at 48 h after cecal ligation and puncture. In contrast, *Kit^W/Wv^* mice started at 8 h with cytokine levels similar to *Cpa3^+/+^* and *Cpa3^Cre/+^* mice, but mounted an almost 10-fold higher TNFα, IL-6 and MCP-1 response at 24 h after CLP (Fig. 1D-F). Sham-treated animals (dashed lines) had transiently increased levels of cytokines at 8 h, in response to the anesthesia and laparotomy (Fig. 1D-F). In Fig. 1G-I we plotted the 24 h serum levels of individual mice, comparing *Cpa3^+/+^, Cpa3^Cre/+^* and *Kit^W/Wv^* mice. Median cytokine levels were similar in *Cpa3^+/+^* and *Cpa3^Cre/+^* mice but elevated in *Kit^W/Wv^* mice. Comparable cytokine responses in *Cpa3^+/+^* and *Cpa3^Cre/+^* mice are consistent with their similar survival rates. Clearly, and in contrast to earlier reports^1,2^, the TNFα response was independent of the presence of mast cells, excluding these cells as the main source for TNFα in experimental peritoneal sepsis. Elevate cytokine responses were associated with the *Kit* mutation. Rather than protection, the strongly elevated inflammatory cytokine levels 24 h after cecal ligation and puncture in most *Kit^W/Wv^*, and in some *Cpa3^+/+^* and *Cpa3^Cre/+^* mice may reflect an exaggerated immune response before succumbing to sepsis.

Collectively, these experiments demonstrate that mast cells do not play a role in protecting against enteric bacterial sepsis in the CLP model. We confirmed the severely impaired survival of *Kit^W/Wv^* mice, and found that this was unrelated to deficiencies in production of TNFα, IL-6, or MCP-1.

### Mast cell-deficient *Kit^W/Wv^* mice resist intraperitoneal injection with cecal bacteria

Kit is essential for the development of interstitial cells of Cajal and for intestinal pacemaker activity, and hence autonomous gut motility is largely abrogated in *Kit* mutants^17,18^. Given this dysfunction of the intestinal physiology in *Kit^W/Wv^* mice, we considered the possibility that the surgical CLP procedure, involving tissue ligation and perforation, could contribute to the sensitivity of *Kit^W/Wv^* mice to sepsis. We therefore induced sepsis without injury of the colon by injection of a cecal bacterial slurry containing a defined amount of enteral bacteria. This is a reproducible, bacteria dose-controlled peritoneal infection model^19^. Donors for enteral slurry were C57BL/6 mice.

More than 90% of all three genotypes (*Kit^W/Wv^*, *Cpa3^+/+^* and *Cpa3^Cre/+^* mice) survived injection of low dose (3 x 10^8^) bacteria (Fig. 2A). Survival was reduced to less than 50% during the observation period of 10 days at an approximately 5-fold higher dose (14 x 10^8^ bacteria) (Fig. 2B). For both doses, the outcome for *Kit^W/Wv^* mice was statistically comparable to that of wild-type control mice (Fig. 2A, B), ruling out immunodeficiency to explain the sensitivity of *Kit^W/Wv^* mice in cecal ligation and puncture. Mast cell-deficient *Cpa3^Cre/+^* mice were no more susceptible than their mast cell bearing littermates (Fig. 2A, B), confirming the irrelevance of mast cells in this bacterial injection model.

**Figure 2.**
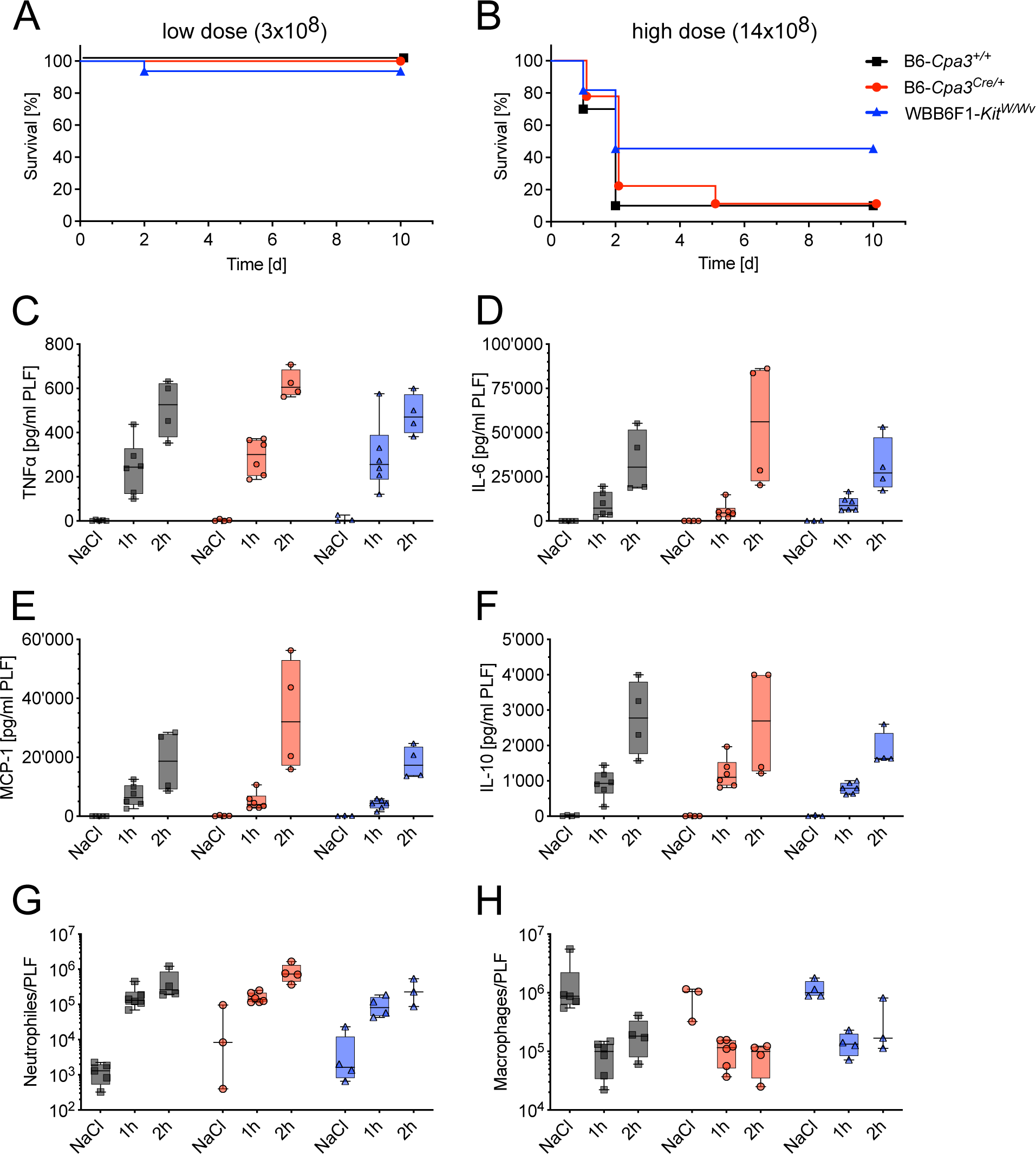
*Kit^W/Wv^* mice are as resistant to injection of cecal slurry as *Kit^+/+^* mice. (A, B). Survival of B6-*Cpa3^+/+^*, B6-*Cpa3^Cre/+^*, and WBB6F1-*Kit^W/Wv^* mice following injection of low dose (3 x 10^8^) bacteria (A), or high dose (14 x 10^8^) bacteria (B) from intestines of normal C57BL/6 mice. For both doses, the outcome for *Kit^W/Wv^* mice was statistically comparable to that of *Kit* wild-type mice, and mast cell deficiency (B6-*Cpa3^+/+^* versus B6-*Cpa3^Cre/+^*) also played no role in survival. Low dose survival: *+/+* 100% of n = 12; *Cre/+* 100% of n = 16; *W/Wv* 94% of n = 16 and p (*+/+* vs *W/Wv*) = 0.3865; high dose survival: *+/+* 10% of n = 10; *Cre/+* 11% of n = 9; *W/Wv* 45% of n = 11 and p (*+/+* vs *W/Wv*) = 0.1110. P values of the survival curve comparisons were calculated using the Mantel-Cox Log-rank test. (C-F). Concentration and kinetic of inflammatory cytokine responses in peritoneal lavage fluid from B6-*Cpa3^+/+^*, B6-*Cpa3^Cre/+^* and WBB6F1-*Kit^W/Wv^* following saline (NaCl) control injection, or one or two hours after low dose bacterial challenge (3-6 x 10^8^ bacteria). Boxes indicate the 25th to 75th percentiles, center lines the median, whiskers the minimum to maximum values, and each dot represents an individual mouse. Statistical analyses were performed on log(Y+1)-transformed values. Ordinary two-way ANOVA with genotype and time as factors revealed no significant effects of genotype or time *x* genotype interaction for any of the cytokines, only significant effects of time were observed (F and P values in Table S1). Sample sizes were n = 6 per genotype at 1 h, n = 4 per genotype at 2 h, and for the NaCl group: n = 5 (*+/+*), n = 4 (*Cre/+*), and n = 3 (*W/Wv*), except for IL-10 where *+/+* mice were n = 4. (G-H). Absolute numbers of (G) neutrophils (Gr1^+^ CD11b^+^) and (H) macrophages (Gr1^−^ CD11b^+^ F4/80^+^) in the peritoneal cavity of mice treated as in (C-F). Statistical analyses were performed on log(Y)-transformed values using ordinary two-way ANOVA with genotype and time as factors. Significant effects of time, but no effects of genotype or time *x* genotype interaction, were observed. (F and P values in Table S1). Sample sizes were for NaCl: n = 5 (*+/+*), n = 3 (*Cre/+*), and n = 4 (*W/Wv*); at 1h: n = 6 (*+/+*), n = 6 (*Cre/+*), and n = 4 (*W/Wv*); and at 2h n = 4 (*+/+*), n = 4 (*Cre/+*), and n = 3 (*W/Wv*).

In order to measure early-stage inflammatory cytokine release, low dose cecal slurry infection was repeated, and mice were analyzed one or two hours later. Compared to NaCl control injected animals, TNFα, IL-6, MCP-1 and IL-10 were elevated in the peritoneal lavage already after one hour, and the cytokine concentrations further increased at two hours after bacteria injection, in particular for IL-6 and MCP-1 (Fig. 2C-F). We could not detect increases in IFN-γ or IL12p70 (data not shown). Statistical comparison of the cytokine responses in *Cpa3^+/+^*, *Cpa3^Cre/+^* and *Kit^W/Wv^* mice revealed a significant effect of time, but no significant effects of genotype or genotype *x* time interaction for the cytokine responses, thus excluding mast cells as a major source of inflammatory cytokines, and indicating that this cytokine response is independent of Kit. Since mast cells have been implicated in the recruitment of innate immune cells, we measured neutrophils and macrophages in the peritoneal lavage fluid by flow cytometry one and two hours after bacterial injection (Fig. 2G+H). Neutrophils which are normally absent, or present only at very low numbers in the peritoneal cavity of naïve mice, were rapidly infiltrating the peritoneal cavity of infected animals (Fig. 2G). Of note, we counted very similar numbers of neutrophils (Gr1^+^ CD11b^+^) in mast cell-proficient *Cpa3^+/+^*, and mast cell-deficient *Cpa3^Cre/+^* mice. Neutrophil recruitment was slightly reduced in *Kit^W/Wv^* mice which could be related to their global Kit-dependent neutropenia^20,21^. Numbers of macrophages (F4/80^+^ CD11b^+^) declined^22^ within the first hour of infection in *Cpa3^+/+^*, *Cpa3^Cre/+^* and *Kit^W/Wv^* mice (Fig. 2H). As for the cytokine responses, no significant effect of genotype was observed for the neutrophil recruitment or macrophage decline. The significant main effect was time.

Taken together, in contrast to the cecal ligation and puncture sepsis results, *Kit^W/Wv^* mice show normal survival, cytokine response and neutrophil recruitment after bacterial slurry injection. This implies that the susceptibility of *Kit^W/Wv^* mice to cecal ligation and puncture sepsis does not reflect defects in immunity, and that immunity is again unaffected by the absence of mast cells. Moreover, our time course experiments show that, within the first hours of peritoneal bacterial infection, mast cells are not major producers of inflammatory cytokines.

### *Kit^W/Wv^* mice harbor intestinal microflora with increased pathogenic potential

In the CLP model, sepsis is driven by the mouse’s own intestinal flora. The selective susceptibility of *Kit^W/Wv^* mice, but not *Cpa3^Cre/+^* mice, in this model raises the question whether impaired intestinal physiology, i.e. hypomotility of gut peristalsis and impaired gastrointestinal transit observed in *Kit*-mutants^18,23^, may influence the composition and pathogenicity of the intestinal microflora. To directly compare the pathogenicity, we isolated enteral bacteria from either *Kit^+/+^* or *Kit^W/Wv^* mice (donor flora) and injected them into recipient cohorts of *Cpa3^+/+^* (Fig. 3A), *Cpa3^Cre/+^* (Fig. 3B), or *Kit^W/Wv^* (Fig. 3C) mice (in this experiment all donor and recipient animals were on the WBB6F1 background). Cecal slurries with five different amounts of bacteria (2, 5, 10, 20, 40 x 10^8^) were applied. This titration experiment revealed a clear correlation between bacterial load and mortality. Of note, we observed a marked leftward shift in the dose-response curves towards lower bacteria concentrations when comparing injections with cecal slurries from *Kit^W/Wv^* and *Kit^+/+^* donors (Fig. 3A-C). Collectively, these data indicate a higher pathogenicity of the microflora harbored in *Kit*-mutants compared to *Kit* wild-type mice.

**Figure 3.**
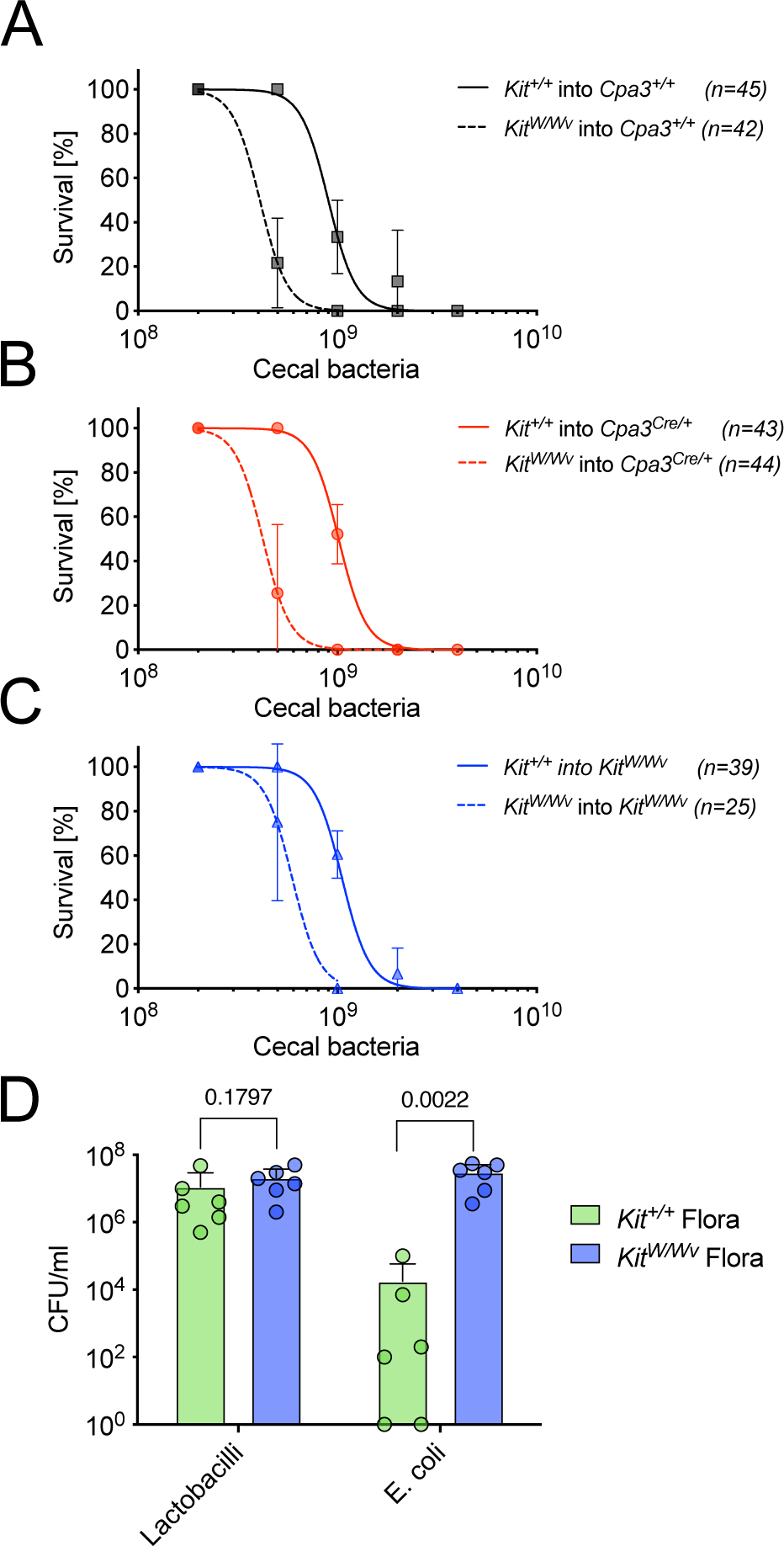
*Kit^W/Wv^* mice harbor more pathogenic enteral bacteria than *Kit^+/+^* mice. (A-C). Survival of *Cpa3^+/+^* (A), *Cpa3^Cre/+^* (B), and *Kit^W/Wv^* (C) mice following injection of increasing doses of bacteria (2, 5, 10, 20 or 40 x 10^8^) isolated from the cecum of *Kit^+/+^* or *Kit^W/Wv^* donor mice. All animals in this experiment were on the WBB6F1 strain background. Group sizes (n) are given in the figure. (D). Bacterial colony forming units were determined for cecal slurries of *Kit^+/+^* (n = 6) and *Kit^W/Wv^* mice (n = 6) which included all cecal slurries used in (A-C). Bars graphs show the mean + SD, each dot represents counts from one individual mouse. P values were calculated with the Mann-Whitney test.

To investigate whether the increased pathogenicity of *Kit^W/Wv^* microbiota is due to an altered bacterial composition of the cecal contents, we first performed a basic microbiological examination and determined bacterial colony forming units (CFU) of the intestinal contents of *Kit^+/+^* and *Kit^W/Wv^* mice, including the specimens used for the i.p. infection experiments shown in Fig. 3A-C. To this end, we prepared serial dilutions of the cecal slurries, cultured these on blood agar and McConkey agar plates, and counted bacterial colonies. While Lactobacilli colonies were obtained at similar frequencies from isolates of both mouse strains, CFU counts for *Escherichia coli (E. coli*) were more that 1000 times higher in *Kit^W/Wv^* than in *Kit^+/+^* isolates (Fig. 3D). *Staphylococcus sp.* (most likely *Staph. Xylosus*) was sporadically found at low level in *Kit^+/+^* slurries (not shown). Hence, *Kit^W/Wv^* microbiota contains high levels of *E. coli*, which may underlie the observed pathogenicity.

### Kit deficiency, but not mast cell deficiency, is associated with intestinal dysbiosis

For a more comprehensive analysis of the intestinal microbiome, we performed 16S rRNA sequencing. The V6 region of the bacterial 16S rRNA gene was amplified and sequenced from cecal samples collected from *Kit^W/Wv^* mice and *Kit^+/+^* littermate controls (Fig. 4A+B), as well as from *Cpa3^Cre/+^* mice and *Cpa3^+/+^* littermate controls (Fig. 4C+D). Amplicon reads were taxonomically classified, and relative abundances were calculated at the order and family levels.

**Figure 4.**
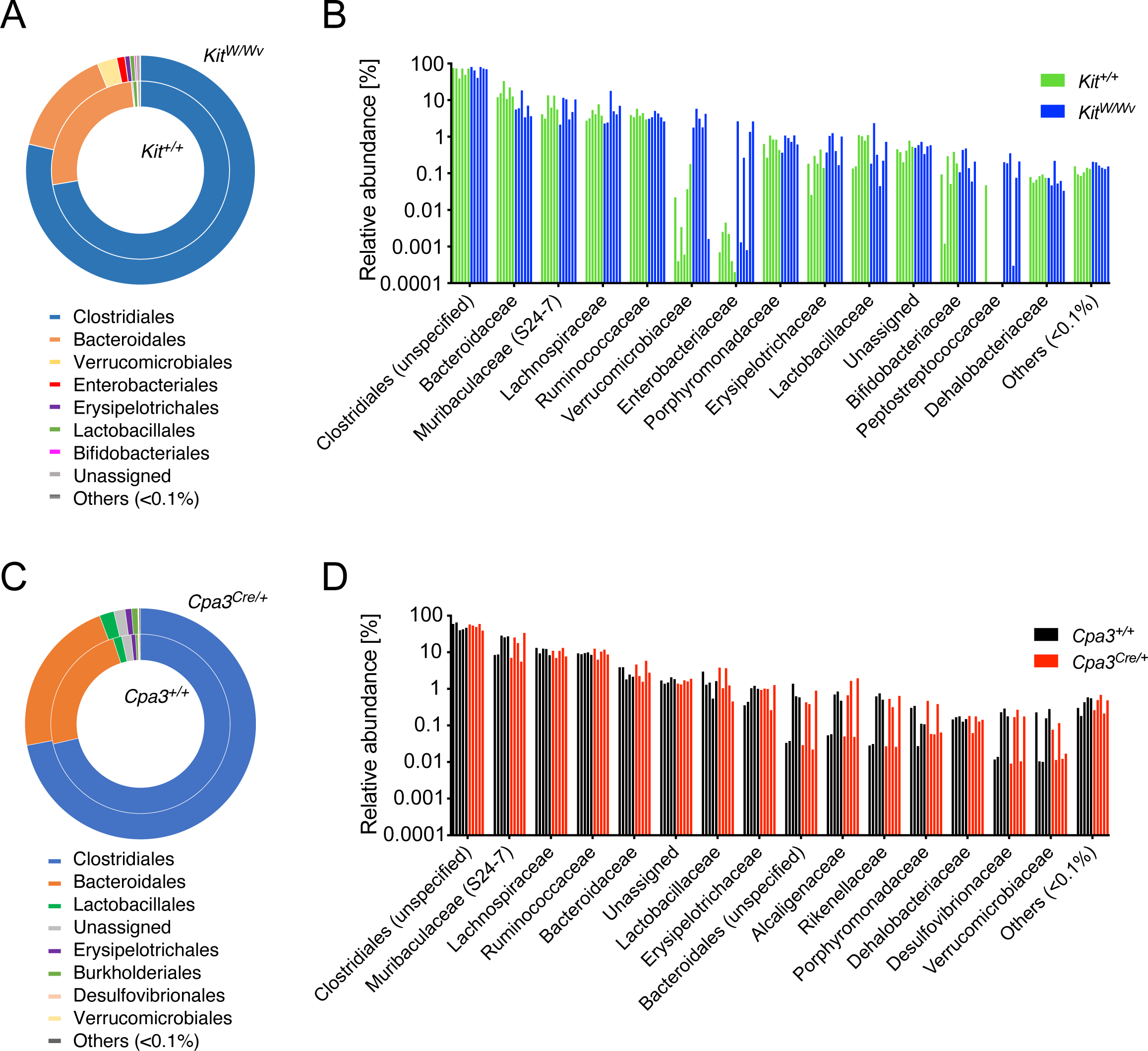
Cecal microbiota composition in *Kit^W/Wv^* and *Cpa3^Cre/+^* mice compared with their respective littermates. (A) Community composition of bacterial orders in cecal samples from *Kit^W/Wv^* (outer circle) and *Kit^+/+^* (inner circle) mice shown as mean relative abundances (n = 6 per group). (B) Relative abundance of bacterial families in individual mice shown as grouped bar plots organized by family. Bars represent data of individual *Kit^+/+^* (green) and *Kit^W/Wv^* (blue) animals. Statistical analysis was performed using ALDeX2. (C) Community composition of bacterial orders in cecal samples from *Cpa3^Cre/+^* (outer circle) and *Cpa3^+/+^* (inner circle) mice shown as mean relative abundances (n = 5 per group). (D) Relative abundance of bacterial families in individual mice shown as grouped bar plots organized by family. Bars represent data of individual *Cpa3^+/+^* (black) and *Cpa3^Cre/+^* (red) animals. Statistical analysis was performed using ALDeX2. Taxa with a relative abundance >0.1% in at least one sample within each panel are shown individually; all other taxa were summed up in “Others”.

At the bacterial order level mean relative abundances revealed similar microbial community structures that were dominated by Clostridiales and Bacteroidales, irrespective of the genotype of the mice (Fig. 4A+C). Nevertheless, host genotype-associated shifts were observed for specific taxa in *Kit^W/Wv^* mice (Fig. 4A outer circle) exhibiting increased relative abundances of Verrucomicrobiales, Enterobacteriales, and Erysipelotrichales, accompanied by a reciprocal decrease in Bacteroidales relative to *Kit^+/+^* controls (inner circle). Lactobacillales and Bifidobacteriales were present at comparable abundances in both genotypes.

To assess host genotype-associated differences in microbiome composition at higher taxonomic resolution, the bacterial family-level read count matrix was analyzed using the statistical package ALDeX2. For visualization, relative abundances were displayed as grouped bar charts depicting individual mice with separate bars (Fig. 4B+D).

Differential abundance analysis found the bacterial families Peptostreptococcaceae, Verrucomicrobiaceae, Enterobacteriaceae, Coriobacteriaceae, and Erysipelotrichaceae enriched in *Kit^W/Wv^* mice compared to *Kit^+/+^* controls, exhibiting moderate-to-large effect sizes (Table S2). In particular, Enterobacteriaceae, Peptostreptococcaceae, and Verrucomicrobiaceae displayed some of the largest positive effect sizes observed in the dataset. Although, these differences did not remain statistically significant after FDR correction, the large differential abundances and effect sizes suggest a biologically relevant restructuring of the microbial community towards facultative anaerobic and potentially pro-inflammatory bacterial families in *Kit^W/Wv^* mice. Moreover, the marked increase in Enterobacteriaceae (diff_abundance = 8.24) in *Kit^W/Wv^* mice, together with the largely unaltered abundance of Lactobacillaceae (effect = -0.54, FDR = 0.939), was consistent with the CFU counts obtained from the cecal slurry titration assay (Fig. 3D). In contrast to these increases, Bacteroidaceae were significantly reduced in *Kit^W/Wv^* mice compared to *Kit^+/+^* controls (effect = −2.40, FDR = 0.018) (Table S2).

Interestingly, in contrast to the pronounced microbiome alterations observed in *Kit^W/Wv^* mice, the microbiome composition of *Cpa3^Cre/+^* mice was largely comparable to that of *Cpa3^+/+^* littermate controls (Fig. 4C+D). ALDEx2 analysis revealed only small effect sizes (effect < 0.7), uniformly high FDR values (FDR > 0.98), and no coherent directional pattern across phylogenetically related bacterial families (Fig. 4D and Table S3).

Collectively, these findings indicate that Kit deficiency, but not mast cell deficiency, is the major determinant of the microbiome alteration observed in *Kit^W/Wv^* mice, and that the increased pathogenicity of their intestinal flora is associated with a dysbiotic shift toward bacterial taxa associated with intestinal inflammation.

### Cohousing transfers sepsis resistance to *Kit^W/Wv^* mice

To obtain further evidence that pathogenic microbiota in *Kit^W/Wv^* mice is responsible for the increased CLP-susceptibility of this strain independent of mast cells, we performed co-housing experiments. *Kit^W/Wv^* and their wild-type *Kit^+/+^* littermates are F1 offspring from WB-*Kit^W/+^* and C57BL/6-*Kit^Wv/+^* parents. The phenotypically different *Kit^W/Wv^* (white) and *Kit^+/+^* (black) mice were normally separated at weaning, which may result in genotype-specific drifts of the enteric microbiota. To prevent such drift, we co-housed *Kit^W/Wv^* and *Kit^+/+^* mice, and repeated cecal ligation and puncture experiments. In contrast to the previous CLP experiments with non-co-housed mice in which severe (22 G needle) conditions were lethal for all *Kit^W/Wv^* mice (Fig. 1B), after co-housing there were no differences in survival (P = 0.3809) comparing *Kit^W/Wv^* (74% of n = 38) and *Kit^+/+^* (65% of n = 25) mice (Fig. 5). These experiments demonstrate that co-housing transfers CLP-resistance from *Kit^+/+^* to *Kit^W/Wv^* mice.

**Figure 5.**
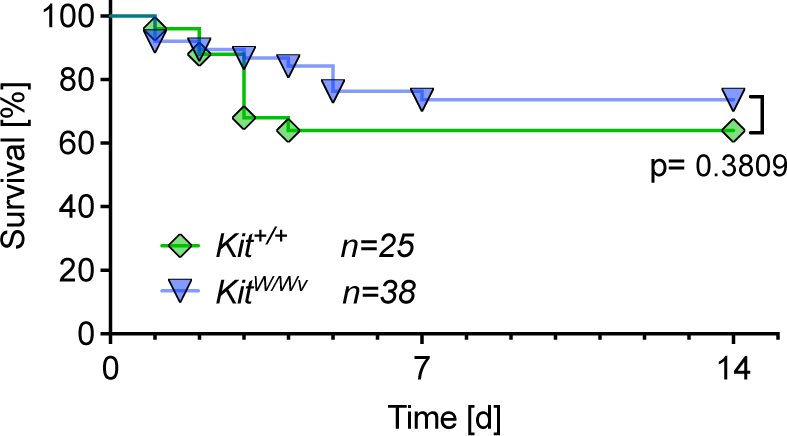
Co-housing of *Kit^W/Wv^* with *Kit^+/+^* mice normalizes the susceptibility of *Kit^W/Wv^* mice to cecal ligation and puncture. *Kit^+/+^* and *Kit^W/Wv^* mice (WBB6F1 background) were housed together from weaning onwards until cecal ligation and puncture (22 G needle). This constant co-housing, increased the survival rate of *Kit^W/Wv^* mice (n=38) and equalized it (p=0.3809) to the probability of survival of *Kit^+/+^* mice (n=25). P values were calculated using the Mantel-Cox Log-rank test.

## Discussion

*Kit^W/Wv^* mice exhibit increased mortality compared to *Kit^+/+^* control mice^1^ in the clinically relevant cecal ligation and puncture (CLP) model of polymicrobial sepsis^14^. This phenotype has been attributed to the absence of mast cells, and broadly been interpreted as evidence for a non-redundant role of mast cells in antibacterial defense during systemic infection. In the present study, we re-examined CLP sepsis experiments using *Cpa3^Cre/+^* mice, a genetic model that separates mast cell deficiency from other pathophysiology in *Kit* mutants. *Cpa3^Cre/+^* mice, which lack mast cells but have intact Kit signaling, showed survival rates comparable to wild-type controls in response to CLP. This finding was highly reproducible across experiments performed over eight years by three independent investigators at two institutions, using large experimental cohorts, supporting the robustness of this observation. Clearly, this large dataset refutes a protective role for mast cells in polymicrobial sepsis elicited by cecal ligation and puncture (CLP).

There are conflicting reports regarding the role of mast cells in the cecal ligation and puncture assay, or in related bacterial infection models. Published results range from pro-pathogenic roles of mast cells^24^ to enhanced vulnerability of mouse mutants with loss of protease genes (Mcpt4; Mcpt6)^16,25^. These reports are difficult to reconcile with our findings demonstrating unimpaired survival of mast cell-deficient (*Cpa3^Cre/+^*) mice. In a peritonitis model induced by injection of a mouse-virulent strain of *Klebsiella pneumoniae*, *Kit^W/Wv^* mice were reported to exhibit impaired bacterial clearance, reduced neutrophil recruitment, and increased expression of TNFα in the peritoneal cavity^2^. In our hands, injecting the ‘natural mixture’ of cecal bacteria for intraperitoneal infections, *Kit^W/Wv^* mice showed comparable survival, TNFα secretion and neutrophil recruitment to the peritoneal cavity, all of which does not agree with the immunodeficiency that Malaviya et al. described^2^. The reasons for these discrepancies are unclear. Naturally, host-to-host transmission of *Klebsiella pneumoniae* requires close contact and generally occurs through the fecal-oral route^26^ rather than experimental intraperitoneal infections. Rather than examining host defense against single bacterial strains, such as *Klebsiella pneumoniae* or *E. coli,* we chose a more natural polymicrobial model, mimicking the release of enteral bacteria following gastrointestinal rupture, and found no evidence for a protective role of mast cells. This conclusion aligns with recent proteome data obtained from primary mouse and human mast cells^27^. By comparison with macrophages, ex vivo isolated mast cells lacked several different classes of pattern recognition receptors which suggests that peritoneal mast cells, at least at steady state, do not possess broad innate pattern recognition capabilities. Specifically, primary human and mouse mast cells do not express detectable amounts of TLRs, and their expression of MYD88 most likely does not represent a configuration for TLR but rather for IL-1R signaling. Mast cells share expression of the RNA sensor RIG-I and its downstream transcription factor IRF3 with most other immune cells, but do not express other RIG-I-like receptors or their downstream signaling components. Mast cells also do not express C-type lectin receptor components, or inflammasome components^27^. Importantly, TNFα protein, reported to be stored in mast cell secretory granules in vitro^28^, was also undetectable in primary mast cells^27^. These data, together with our polymicrobial sepsis experiments, make it likely that mast cells play no role in protection against peritoneal sepsis. Instead, it appears that the principal innate responder cells in peritoneal sepsis are macrophages and neutrophils which are also the main cellular sources of TNFα.

How is it possible that the opposite conclusion, i.e. that mast cells are key in protection from sepsis, had been reached 30 years ago based on experiments in *Kit* mutants? Or, in other words, if their mast cell deficiency is not causal for sepsis susceptibility, what renders *Kit* mutants so susceptible? We also addressed this question. The answer came from focusing in particular on intestinal flora. When we challenged *Kit^W/Wv^* mice, which are also in our hands sensitive to CLP sepsis, through intraperitoneal injection of intestinal bacterial suspensions (“cecal slurry”) from wild type intestines, *Kit^W/Wv^* mice were as resistant as normal mice to the infection. Of note, *Kit^W/Wv^* mice exhibited survival kinetics, cytokine responses, and neutrophil recruitment indistinguishable to *Cpa3^Cre/+^* mice or *Cpa3^+/+^* controls. Interestingly, we next discovered that cecal slurries prepared from *Kit^W/Wv^* intestines were more pathogenic and contained substantially higher *E. coli* CFU than those from wild-type controls. In brief, the increased susceptibility of *Kit*-mutant mice to CLP is unrelated to mast cells or other defects in antibacterial immunity, but instead reflects differences in the infectious inoculum released upon intestinal injury.

Consistent with the impaired survival and *E. coli* CFU data, 16S rRNA sequencing of cecal samples revealed an altered microbial ecosystem in *Kit^W/Wv^* mice compared to *Kit^+/+^* controls. *Kit*-mutant mice exhibited a compositional shift characterized by an enrichment of Enterobacteriaceae, Verrucomicrobiaceae, Erysipelotrichaceae, and Peptostreptococcaceae, along with a reduction in Bacteroidaceae. Members of the Enterobacteriaceae family, including Escherichia coli, are facultatively anaerobic Gram-negative bacteria with a well-established pathogenic potential in sepsis. They are frequent drivers of inflammation following intestinal barrier disruption^29,30^. Likewise, Erysipelotrichaceae and Peptostreptococcaceae have been associated with intestinal inflammation, inflammatory bowel disease, and microbiota instability in both humans and experimental models^29,31^. Notably, the increase of Erysipelotrichaceae observed in *Kit^W/Wv^* mice was mainly attributed to the genus *Allobaculum*. Recent studies identified *Allobaculum mucolyticum* as a mucin-degrading bacterium in the intestines of IBD patients, suggesting that an expansion of this genus may reflect altered ecological conditions in the mucosa during inflammatory diseases^32^. A parallel expansion of Peptostreptococcaceae and Enterobacteriaceae, together with a concomitant decrease of Bacteroidaceae, has been reported for example in colitis and IBD patients^33,34^. Collectively, the microbial changes we observed in *Kit^W/Wv^* mice resemble dysbiotic patterns reported in chronic intestinal inflammation, experimental colitis, and impaired barrier function. We note that the observed microbiome differences are correlative and do not formally demonstrate which specific bacterial taxa were responsible for the increased pathogenicity observed in the CLP or cecal slurry assays.

Given that *Cpa3^Cre/+^* mice relative to *Cpa3^+/+^* controls have an unaltered intestinal microbiome, we exclude mast cell deficiency as the causal factor for the underlying dysbiosis in *Kit^W/Wv^* mice. *Kit*-mutant mice suffer from a known intestinal motility disorder, resulting from impaired development and maintenance of interstitial cells of Cajal^17,18^. While it was not a focus of this work to directly address the relationship between intestinal motility and microbial composition, the microbiota alterations identified by 16S rRNA gene sequencing are compatible with delayed intestinal transit reshaping the gut microbial ecosystem. This condition may favor the expansion of facultative anaerobic bacteria such as Enterobacteriaceae, and thus contribute to a dysbiotic microbial community associated with intestinal inflammation.

The differences in composition and pathogenicity of *Kit-*mutant and wild-type cecal microbiota we report here can only directly refer to the mice bred and maintained during the experimental period in our mouse facilities. Breeding *Kit^+/+^* and *Kit^W/Wv^* mice from WB*-Kit^W/+^* and B6-*Kit^Wv/+^* parents, and separating the offspring at the time of weaning, led to the observed differences in microbiota within the same animal facility. Performing sepsis experiments in mice generated in this manner revealed that lack of Kit can strongly enhance the pathogenicity and, in particular, markedly increase numbers of *E. coli* colony-forming units. We suggest assessing beforehand whether microbiota are, in fact, comparable between wild-type and mutant mice in CLP sepsis experiments. If they are not, as in the case of the *Kit* mutants, the experimental and the control group may release incomparable microbiota from the intestinal puncture. This consideration also applies to experimental and control mice obtained from different breeders or housed separately.

Collectively, our experiments show that mast cells play no role in protection from endomicrobial sepsis, and they also provide an explanation for the apparent immunological susceptibility and mast cell-dependent phenotype of *Kit* mutants in this infection.

## Supporting information

Supplemental Items

## Acknowledgements

We thank Andrea Erles-Kemna, Katja Schmidt and Werner Nicklas, DKFZ, for expert microbiological analyses, and the staff of the animal facilities at the University Clinics Ulm and at DKFZ for expert mouse husbandry. We thank Bernd Echtenacher for the practical introduction to the CLP procedure, and Sven Schäfer for help with CLP experiments. We are grateful to Konrad Bode for his expertise in microbiome analysis. We thank the NGS Core Facility, DKFZ, for expert sequencing. We thank Axel Roers and Thomas Plum for critical reading of the manuscript. This work was supported by the Deutsche Forschungsgemeinschaft (DFG) CRC156/TRR156 project A7 to T.B.F. and H.-R.R., and ERC Advanced grant 233074, HGF Project Immunology & Inflammation (ZT-0027), and the Leibniz program of the DFG (all to H.-R.R.).

## Methods

### Mice

B6-*Cpa3^Cre^*^/+^ mice were obtained after backcrossing the *Cpa3^Cre^* allele (*Cpa3^tm3(icre)Hrr^*)^6^ for >20 generations onto C57BL/6J (B6). In all experiments, littermates from the breeding of B6-*Cpa3^Cre^*^/+^ with wild-type B6 mice were used. *Kit-*mutant mast-cell-deficient WBB6F1*-Kit^W/Wv^* mice and their WBB6F1*-Kit*^+/+^ littermate controls were obtained by crossing WB-*Kit^W^*^/+^ with B6-*Kit^Wv^*^/+^ mice (both originally obtained from Japan-SLC, Shizuoka, Japan). Unless otherwise described in the text, WBB6F1*-Kit^W/Wv^* mice and their WBB6F1*-Kit*^+/+^ littermates were separated by coat color at weaning. For direct comparison, WBB6F1-*Cpa3^Cre^*^/+^ and WBB6F1-*Cpa3*^+/+^ littermates were generated in F1 intercrosses of WB-*Kit*^+/+^ with B6-*Cpa3^Cre^*^/+^ mice.

All animals were generated at the mouse breeding facilities of the University Clinics in Ulm or the center for preclinical research at the DKFZ in Heidelberg. All experiments were conducted in accordance to animal care guidelines pertaining to local animal committees (Regierungspräsidium Tübingen or Regierungspräsidium Karlsruhe) and to the institutional guidelines.

### Sepsis models

#### Cecal ligation and puncture (CLP)

The surgical sepsis model was performed according to Rittirsch et al.^35^ and Cuenca et al.^36^. In brief, mice were anesthetized (100 mg/kg ketamine and 16 mg/kg xylazine in saline, i.p.), belly shaved and placed onto a sterile-covered warming plate for the duration of the surgery. After disinfection with 70% ethanol, the abdomen of the mice was opened with scissors by a 1-cm mid line cut into the skin, followed by a 1-cm incision of the peritoneum. The cecum was exteriorized and its stool content was gently palpated towards the distal end. The ligation with 4-0 sutures was placed at half the distance (‘50% ligation’) between the distal pole and the base of the cecum. A single puncture was applied to the pole of the cecum (gauge sizes of the used needles are indicated for each experiment). Gentle pressure on the ligated cecum released a tiny drop of the cecal content while the needle was withdrawn to prevent immediate closure of the puncture hole. Afterwards the cecum was reposed and wound closure was performed with surgical clips. Sham control mice underwent the same surgical procedure, however without ligation and puncture of the cecum. Finally, mice received 1 ml saline s.c. for fluid replacement. For the entire observation period after surgery, all mice were closely monitored, and animals showing signs of severe distress or suffering were euthanized. Hence, statements in this manuscript referring to mortality all fell under this animal welfare protocol. For profiling of inflammatory cytokines, mice were sacrificed at the indicated time after surgery and blood for serum preparation was collected by heart puncture.

#### Cecal slurry injection

For the non-surgical sepsis model, suspensions of cecal bacteria were prepared from donor mice and injected intraperitoneally into the test mice. For preparation of the cecal slurry, four donor mice (B6-*Cpa3^+/+^* (Fig. 2), WBB6F1-*Kit^+/+^* or WBB6F1-*Kit^W/Wv^* (Fig. 3)) were sacrificed and the content of their cecum was pooled and resuspended in 10 ml saline (0.9 M NaCl). The suspension was centrifuged for 1 min at 20 g (200 rpm) in a swing-out bucket and the upper 8 ml were subsequently passed through 100 μm and 40 μm filters. Bacteria counts (rod-shaped) were determined using a Neubauer counting chamber with 0.02 mm chamber depth. Injected bacterial doses are indicated in Figures 2 and 3. Injected mice were closely monitored and moribund mice were sacrificed (see ‘Cecal ligation and puncture’ paragraph above). For the analysis of cytokine concentrations and myeloid cell infiltrations in the peritoneal cavity, mice were sacrificed at the indicated time after cecal slurry injection and the peritoneal fluid was collected. Therefore, 700-800 μl sterile FACS buffer (PBS with 5% FCS) were injected intraperitoneally, and after a short abdominal massage, the peritoneal cavity was opened by a 5 mm incision to retrieve about 500 μl of the peritoneal exudate suspension with a pipette. A small aliquot was used to determine the concentration of cells and the total amount of cells was calculated from the total injected volume. The peritoneal exudate suspension was then separated into a cellular and a soluble fraction by centrifugation for 5 min with 500 g (2200 rpm). The cell pellet was resuspended in FACS buffer for subsequent flow cytometric analysis. The soluble fraction was centrifuged for another 5 min at 16200 g (13000 rpm) and the supernatant was snap frozen in liquid nitrogen for later cytokine measurement.

#### CFU determination

CFU of the cecal slurry suspensions used in the sepsis experiment was determined by plating serial dilutions (10^-3^ – 10^-10^ in 0.9 M NaCl) of the slurry stock suspension in replicates on blood agar and McConkey agar. *Lactobacillus* and *Staphylococcus* colonies were counted on blood agar plates, *Escherichia coli* colonies on McConkey agar plates.

### Microbiome analysis

Mice were housed in individually ventilated cages under barrier conditions at the central animal facility of the German Cancer Research Center, Heidelberg, and fed standard breeding chow (Kliba 3307). After weaning, littermates were separated and maintained in individual cages for at least nine weeks before sacrifice and cecal content collection. DNA was extracted using the QIAamp DNA Stool Mini Kit (Qiagen). The V6 region of the bacterial 16S rRNA gene was amplified using indexed primers (see Table S4) under the following conditions: 94 °C for 3 min; 32 cycles of 94 °C for 45 s, 52 °C for 60 s, and 72 °C for 90 s; followed by 72 °C for 10 min. PCR amplicons were purified using the QIAquick PCR Purification Kit (Qiagen) and equimolarly pooled for library preparation at the NGS core facility. Paired-end sequencing was performed on the Illumina MiSeq platform (Illumina, San Diego, CA, USA).

Raw reads were quality-trimmed and denoised using DADA2 (R v3.6.3). Resulting amplicon sequence variants were taxonomically classified at phylum, class, order, family, and genus levels using the SILVA database (v132). Sequencing depth ranged from 90k to 850k reads per sample.

Differential abundance analysis at the family level was performed in R (v4.6.0) using ALDEx2 (v1.44.0) on raw count data. Taxa were filtered by prevalence (≥20% of samples) and total abundance (>5 reads across all samples) prior to analysis. ALDEx2 used 128 Monte Carlo Dirichlet instances and centered log-ratio-transformed counts to estimate effect sizes, differential abundance, and Welch’s t-test P values with Benjamini–Hochberg FDR correction.

Relative abundances profiles were calculated by normalizing raw read counts to total sequencing depth per sample. In graphical illustrations, low-abundance taxa with a mean relative abundance <0.1% were collapsed into a single “Others (<0.1%)” category.

### Flow cytometric assays

#### Cytometric bead array (CBA)

Cytokine concentrations of serum samples or peritoneal lavages were determined in a bead-based multiplex cytometric assay (Cytometric Bead Array (CBA) Mouse Inflammation Kit, BD Biosciences). Samples (undiluted and 1:20 dilutions) and cytokine standards (10 ng – 5 pg serial dilutions) were incubated with mixtures of fluorescent beads coated with capture antibodies against IL-6, IL-10, INF-γ, MCP-1, TNF-α and IL12p70. PE-labeled detection reagent (i.e. PE-conjugated antibodies specific for the respective mouse cytokines) was added, and, after incubation and wash, samples were measured at a BD FACSCanto^TM^ or BD LSRFortessa^TM^ instrument. Cytokine concentrations were calculated from mean fluorescence values using the BD FCAP Array Software.

#### Peritoneal exudate cells (PEC)

The cellular fraction of the peritoneal exudate suspensions was analyzed for CD117^+^ mast cells, Gr1^+^ CD11b^+^ granulocytes and Gr1^−^ CD11b^+^ F4/80^+^ macrophages on a BD FACS Canto. Therefore, cells were first incubated with mouse IgG (300 μg/ml, Dianova) for blocking of Fc-receptors and then stained with titrated amounts of fluorescent-labeled antibodies in PBS with 5% FCS. The following antibodies were used: CD117-APC (2B8), CD11b-FITC (M1/70), Gr1-PE (RB6-8C5) all from Pharmingen, and F4/80-APC-A780 (BM8) from eBioscience.

