## Supplemental Items for "Role of mast cells in endomicrobial sepsis revisited: Mast cell-deficient mice show normal immunological protection"

Table S1. Statistical summary of Fig. 2C-H

Table S2. Differential abundance analysis of cecal bacterial families in $Kit^{W/Wv}$ and $Kit^{+/+}$ mice using ALDEx2.

Table S3. Differential abundance analysis of cecal bacterial families in $Cpa3^{Cre/+}$ and $Cpa3^{+/+}$ mice using ALDEx2.

Table S4: Primers used for the next generation sequencing of the V6 region of the 16S rRNA gene

**Table S1. Statistical summary of Fig. 2C-H.**

Results of ordinary two-way ANOVA performed on log(Y+1)-transformed data for panels C–F and log(Y)-transformed data for panels G+H. Time and genotype were included as independent factors, and the interaction term represents the combined effect of time and genotype. F values are presented as F($df_1$, $df_2$), where $df_1$ denotes the degrees of freedom for the factor and $df_2$ denotes the residual degrees of freedom, together with corresponding P values.

| Figure panels | Time | | Genotype | | Time *x* Genotype Interaction | |
|---|---|---|---|---|---|---|
| C | F (2, 33) = 183.9 | P<0.0001 | F (2, 33) = 0.5753 | P=0.5681 | F (4, 33) = 0.4711 | P=0.7565 |
| D | F (2, 33) = 413.9 | P<0.0001 | F (2, 33) = 0.5637 | P=0.5745 | F (4, 33) = 1.787 | P=0.1551 |
| E | F (2, 33) = 129.1 | P<0.0001 | F (2, 33) = 0.02416 | P=0.9761 | F (4, 33) = 0.5720 | P=0.6848 |
| F | F (2, 32) = 129.5 | P<0.0001 | F (2, 32) = 0.3138 | P=0.7329 | F (4, 32) = 0.1849 | P=0.9446 |
| G | F (2, 30) = 70.47 | P<0.0001 | F (2, 30) = 3.023 | P=0.0637 | F (4, 30) = 1.106 | P=0.3718 |
| H | F (2, 30) = 35.67 | P<0.0001 | F (2, 30) = 2.335 | P=0.1142 | F (4, 30) = 0.8764 | P=0.4896 |

**Table S2. Differential abundance analysis of cecal bacterial families in *Kit*$^{W/Wv}$ and *Kit*$^{+/+}$ mice using ALDEx2.**

Family-level differential abundance assessed by ALDEx2 from 16S rRNA sequencing read counts. Shown are effect size (effect), differential abundance between groups (diff_abundance), Welch's t-test P value (p_value), Benjamini–Hochberg false discovery rate-adjusted P value (fdr), and direction of change. Taxa are ranked according to the absolute effect size.

| taxa | effect | diff_abundance | p_value | fdr | direction |
|---|---|---|---|---|---|
| f__Bacteroidaceae | -2.4013 | -1.8037 | 0.0004 | 0.0184 | Higher in WT |
| f__Peptostreptococcaceae | 1.3682 | 9.2868 | 0.0082 | 0.3077 | Higher in W/Wv |
| f__Verrucomicrobiaceae | 1.2542 | 6.8519 | 0.0183 | 0.3857 | Higher in W/Wv |
| f__Enterobacteriaceae | 1.0158 | 8.2389 | 0.0312 | 0.4607 | Higher in W/Wv |
| f__Deferribacteraceae | -0.9894 | -5.4973 | 0.0467 | 0.6286 | Higher in WT |
| f__Coriobacteriaceae | 0.9281 | 1.2133 | 0.0618 | 0.6128 | Higher in W/Wv |
| f__Ruminococcaceae | -0.7939 | -0.6420 | 0.1758 | 0.8171 | Higher in WT |
| f__Erysipelotrichaceae | 0.6729 | 1.4534 | 0.1087 | 0.8426 | Higher in W/Wv |
| f__Paenibacillaceae | -0.6153 | -2.3802 | 0.1903 | 0.7445 | Higher in WT |
| f__Dehalobacteriaceae | -0.5785 | -0.6018 | 0.2004 | 0.8358 | Higher in WT |
| f__Streptococcaceae | -0.5609 | -1.8035 | 0.2702 | 0.8204 | Higher in WT |
| f__Lactobacillaceae | -0.5424 | -1.0128 | 0.2775 | 0.9386 | Higher in WT |
| f__Turicibacteraceae | 0.4843 | 2.6701 | 0.3468 | 0.9340 | Higher in W/Wv |
| f__Acholeplasmataceae | -0.4635 | -2.2325 | 0.3596 | 0.8826 | Higher in WT |
| f__Prevotellaceae | -0.4635 | -1.7878 | 0.3622 | 0.8723 | Higher in WT |
| f__Muribaculaceae (S24-7) | -0.4379 | -0.6128 | 0.3673 | 0.9483 | Higher in WT |
| o__RF32 (unspecified) | -0.4335 | -1.9219 | 0.3088 | 0.8220 | Higher in WT |
| f__Burkholderiaceae | -0.4153 | -1.7025 | 0.3695 | 0.8486 | Higher in WT |
| f__[Mogibacteriaceae] | -0.3658 | -0.4689 | 0.3552 | 0.9219 | Higher in WT |
| f__Bacillaceae | -0.3586 | -1.6257 | 0.4560 | 0.9197 | Higher in WT |
| f__Planococcaceae | -0.3355 | -1.5388 | 0.4213 | 0.8938 | Higher in WT |
| o__Lactobacillales (unspecified) | -0.3258 | -1.5326 | 0.5094 | 0.9279 | Higher in WT |
| f__[Odoribacteraceae] | -0.3023 | -1.5216 | 0.5524 | 0.9393 | Higher in WT |
| o__Clostridiales (unspecified) | -0.2853 | -0.3546 | 0.5795 | 0.9847 | Higher in WT |
| f__Pasteurellaceae | 0.2614 | 1.4195 | 0.5983 | 0.9610 | Higher in W/Wv |
| f__Carnobacteriaceae | -0.2164 | -1.0390 | 0.5490 | 0.9370 | Higher in WT |
| Unassigned | -0.2134 | -0.2806 | 0.8036 | 0.9927 | Higher in WT |
| f__Desulfovibrionaceae | 0.1902 | 0.7063 | 0.5783 | 0.9890 | Higher in W/Wv |
| f__Peptococcaceae | -0.1878 | -0.7323 | 0.6429 | 0.9588 | Higher in WT |
| f__Pseudomonadaceae | -0.1845 | -0.8488 | 0.6604 | 0.9522 | Higher in WT |
| f__Rikenellaceae | 0.1820 | 0.7928 | 0.6335 | 1.0000 | Higher in W/Wv |
| f__Aerococcaceae | -0.1571 | -0.7634 | 0.6300 | 0.9633 | Higher in WT |
| f__Oleiphilaceae | 0.1450 | 0.6234 | 0.5942 | 0.9631 | Higher in W/Wv |
| o__RF39 (unspecified) | 0.1402 | 0.5705 | 0.6738 | 0.9999 | Higher in W/Wv |
| f__Staphylococcaceae | 0.1318 | 0.6296 | 0.6698 | 0.9794 | Higher in W/Wv |
| f__Propionibacteriaceae | -0.1046 | -0.4793 | 0.6690 | 0.9781 | Higher in WT |
| f__Corynebacteriaceae | -0.1024 | -0.5341 | 0.6330 | 0.9713 | Higher in WT |
| o__Bacteroidales (unspecified) | 0.1020 | 0.4100 | 0.7105 | 1.0000 | Higher in W/Wv |
| f__Alcaligenaceae | -0.0945 | -0.2506 | 0.8026 | 0.9969 | Higher in WT |
| f__Conexibacteraceae | 0.0900 | 0.5027 | 0.7047 | 0.9832 | Higher in W/Wv |
| f__Bradyrhizobiaceae | 0.0807 | 0.3567 | 0.6984 | 0.9805 | Higher in W/Wv |
| f__Lachnospiraceae | -0.0727 | -0.1200 | 0.7481 | 0.9958 | Higher in WT |
| f__Micrococcaceae | -0.0609 | -0.2593 | 0.7058 | 0.9713 | Higher in WT |
| f__Christensenellaceae | -0.0597 | -0.3292 | 0.6944 | 0.9555 | Higher in WT |
| f__Moraxellaceae | -0.0520 | -0.1169 | 0.8328 | 0.9991 | Higher in WT |
| f__Bifidobacteriaceae | 0.0390 | 0.0968 | 0.5058 | 1.0000 | Higher in W/Wv |
| f__Porphyromonadaceae | -0.0388 | -0.0624 | 0.7774 | 0.9957 | Higher in WT |
| f__Comamonadaceae | -0.0227 | -0.0790 | 0.7789 | 0.9935 | Higher in WT |
| f__Clostridiaceae | -0.0167 | -0.0461 | 0.8052 | 1.0000 | Higher in WT |
| f__Xanthomonadaceae | 0.0111 | 0.0867 | 0.7069 | 0.9870 | Higher in W/Wv |

**Table S3. Differential abundance analysis of cecal bacterial families in *Cpa3*$^{Cre/+}$ and *Cpa3*$^{+/+}$ mice using ALDEx2.**

Family-level differential abundance assessed by ALDEx2 from 16S rRNA sequencing read counts. Shown are effect size (effect), differential abundance between groups (diff_abundance), Welch's t-test P value (p_value), Benjamini–Hochberg false discovery rate-adjusted P value (fdr), and direction of change. Taxa are ranked according to the absolute effect size.

| taxa | effect | diff_abundance | p_value | fdr | direction |
|---|---|---|---|---|---|
| f__Planococcaceae | 0.6914 | 1.2267 | 0.1787 | 0.9956 | Higher in Cre |
| f__Turicibacteraceae | 0.6729 | 1.5597 | 0.1318 | 0.9993 | Higher in Cre |
| f__Staphylococcaceae | -0.6585 | -1.2523 | 0.1729 | 0.9990 | Higher in WT |
| f__Bifidobacteriaceae | 0.6279 | 0.4586 | 0.1685 | 0.9998 | Higher in Cre |
| f__SBYG_4172 | -0.4186 | -1.3005 | 0.3005 | 1.0000 | Higher in WT |
| f__Micrococcaceae | -0.4052 | -2.4341 | 0.3842 | 0.9978 | Higher in WT |
| f__Aerococcaceae | 0.3938 | 1.5299 | 0.4057 | 0.9991 | Higher in Cre |
| f__Verrucomicrobiaceae | -0.3233 | -0.7702 | 0.4578 | 1.0000 | Higher in WT |
| o__YS2 (unspecified) | -0.3008 | -1.2553 | 0.3517 | 1.0000 | Higher in WT |
| f__Bacillaceae | -0.2924 | -1.0484 | 0.4708 | 0.9993 | Higher in WT |
| f__Erysipelotrichaceae | 0.2894 | 0.2957 | 0.4731 | 1.0000 | Higher in Cre |
| f__Alcaligenaceae | 0.2738 | 0.8083 | 0.5677 | 1.0000 | Higher in Cre |
| o__RF32 (unspecified) | -0.2616 | -0.7215 | 0.6038 | 1.0000 | Higher in WT |
| f__Bradyrhizobiaceae | 0.2495 | 1.0195 | 0.4953 | 0.9952 | Higher in Cre |
| o__Clostridiales (unspecified) | 0.2391 | 0.2724 | 0.6615 | 1.0000 | Higher in Cre |
| f__Conexibacteraceae | 0.2386 | 1.0700 | 0.5686 | 0.9820 | Higher in Cre |
| f__Comamonadaceae | -0.2324 | -0.8771 | 0.7488 | 1.0000 | Higher in WT |
| f__Propionibacteriaceae | 0.2261 | 0.8119 | 0.6069 | 1.0000 | Higher in Cre |
| f__Deferribacteraceae | -0.2256 | -0.2447 | 0.5747 | 1.0000 | Higher in WT |
| f__Clostridiaceae | 0.2246 | 0.1574 | 0.7041 | 1.0000 | Higher in Cre |
| f__Ruminococcaceae | 0.1965 | 0.1933 | 0.5908 | 1.0000 | Higher in Cre |
| f__Burkholderiaceae | 0.1885 | 0.7994 | 0.5616 | 1.0000 | Higher in Cre |
| f__Bacteroidaceae | 0.1857 | 0.3410 | 0.6306 | 1.0000 | Higher in Cre |
| o__RF39 (unspecified) | 0.1808 | 0.3896 | 0.6289 | 1.0000 | Higher in Cre |
| f__Moraxellaceae | -0.1771 | -0.2545 | 0.7146 | 1.0000 | Higher in WT |
| f__Paenibacillaceae | 0.1642 | 0.3338 | 0.5946 | 1.0000 | Higher in Cre |
| f__Streptococcaceae | 0.1603 | 0.2599 | 0.6946 | 1.0000 | Higher in Cre |
| f__Peptostreptococcaceae | -0.1344 | -0.3519 | 0.8123 | 1.0000 | Higher in WT |
| f__Lactobacillaceae | 0.1279 | 0.2142 | 0.7050 | 1.0000 | Higher in Cre |
| f__ Muribaculaceae (S24-7) | -0.1260 | -0.1119 | 0.8601 | 1.0000 | Higher in WT |
| f__Enterobacteriaceae | 0.1237 | 0.3748 | 0.7890 | 1.0000 | Higher in Cre |
| f__Xanthomonadaceae | -0.1186 | -0.5158 | 0.6811 | 1.0000 | Higher in WT |
| o__Bacteroidales (unspecified) | -0.1120 | -0.3376 | 0.7919 | 1.0000 | Higher in WT |
| f__Prevotellaceae | 0.1113 | 0.3617 | 0.9045 | 1.0000 | Higher in Cre |
| f__Microbacteriaceae | -0.1110 | -0.4765 | 0.6596 | 1.0000 | Higher in WT |
| f__Corynebacteriaceae | 0.1035 | 0.4274 | 0.7458 | 1.0000 | Higher in Cre |
| f__[Mogibacteriaceae] | 0.1022 | 0.1608 | 0.7146 | 1.0000 | Higher in Cre |
| o__Lactobacillales (unspecified) | -0.0982 | -0.3272 | 0.6507 | 1.0000 | Higher in WT |
| f__Acholeplasmataceae | 0.0957 | 0.3579 | 0.8491 | 1.0000 | Higher in Cre |
| f__Lachnospiraceae | 0.0928 | 0.0944 | 0.9188 | 1.0000 | Higher in Cre |
| f__Pasteurellaceae | 0.0925 | 0.3582 | 0.6949 | 0.9963 | Higher in Cre |
| Unassigned | 0.0902 | 0.0590 | 0.7473 | 1.0000 | Higher in Cre |
| f__Christensenellaceae | 0.0764 | 0.2093 | 0.8694 | 1.0000 | Higher in Cre |
| f__Pseudomonadaceae | 0.0756 | 0.2592 | 0.6688 | 1.0000 | Higher in Cre |
| f__Thermoactinomycetaceae | 0.0645 | 0.1371 | 0.7639 | 0.9941 | Higher in Cre |
| f__Oleiphilaceae | -0.0596 | -0.3627 | 0.7135 | 0.9978 | Higher in WT |
| f__Desulfovibrionaceae | -0.0528 | -0.1274 | 0.9394 | 1.0000 | Higher in WT |
| f__Porphyromonadaceae | 0.0415 | 0.1025 | 0.8542 | 1.0000 | Higher in Cre |
| f__[Odoribacteraceae] | -0.0279 | -0.0911 | 0.8589 | 1.0000 | Higher in WT |
| f__Rikenellaceae | -0.0270 | -0.0966 | 0.9353 | 1.0000 | Higher in WT |
| f__Carnobacteriaceae | -0.0262 | -0.1604 | 0.8244 | 1.0000 | Higher in WT |
| f__Peptococcaceae | -0.0253 | -0.0747 | 0.9370 | 1.0000 | Higher in WT |
| f__Dehalobacteriaceae | -0.0157 | -0.0208 | 0.8968 | 1.0000 | Higher in WT |
| f__Coriobacteriaceae | 0.0063 | 0.0130 | 0.8422 | 1.0000 | Higher in Cre |

**Table S4: Primers used for the next generation sequencing of the V6 region of the 16S rRNA gene.**

A) The lower-case part indicates the index sequences used to demultiplex the individual samples after sequencing.

| Mouse ID | Primer forward | Primer reverse |
|---|---|---|
| S650 | gcagt CAACGCGARGAACCTTACC | actgc ACAACACGAGCTGACGAC |
| S672 | tgca CAACGCGARGAACCTTACC | tgca ACAACACGAGCTGACGAC |
| S656 | cgtcga CAACGCGARGAACCTTACC | tcgacg ACAACACGAGCTGACGAC |
| S666 | tgac CAACGCGARGAACCTTACC | gtca ACAACACGAGCTGACGAC |
| S667 | agta CAACGCGARGAACCTTACC | tact ACAACACGAGCTGACGAC |
| S658 | gtcgc CAACGCGARGAACCTTACC | gcgac ACAACACGAGCTGACGAC |
| S654 | tagct CAACGCGARGAACCTTACC | agcta ACAACACGAGCTGACGAC |
| S664 | acgta CAACGCGARGAACCTTACC | tacgt ACAACACGAGCTGACGAC |
| S642 | catgcg CAACGCGARGAACCTTACC | cgcatg ACAACACGAGCTGACGAC |
| S670 | atga CAACGCGARGAACCTTACC | tcat ACAACACGAGCTGACGAC |
| S655 | gactgt CAACGCGARGAACCTTACC | acagtc ACAACACGAGCTGACGAC |
| S665 | cactac CAACGCGARGAACCTTACC | gtagtg ACAACACGAGCTGACGAC |
| S18367 | cactac CAACGCGARGAACCTTACC | agt ACAACACGAGCTGACGAC |
| S18368 | tgac CAACGCGARGAACCTTACC | agt ACAACACGAGCTGACGAC |
| S18189 | gactgt CAACGCGARGAACCTTACC | agt ACAACACGAGCTGACGAC |
| S18166 | tagct CAACGCGARGAACCTTACC | agt ACAACACGAGCTGACGAC |
| S18163 | act CAACGCGARGAACCTTACC | agt ACAACACGAGCTGACGAC |
| S18366 | acgta CAACGCGARGAACCTTACC | agt ACAACACGAGCTGACGAC |
| S18190 | cgtcga CAACGCGARGAACCTTACC | agt ACAACACGAGCTGACGAC |
| S18164 | catgcg CAACGCGARGAACCTTACC | agt ACAACACGAGCTGACGAC |
| S18365 | gtcgc CAACGCGARGAACCTTACC | agt ACAACACGAGCTGACGAC |
| S18165 | gcagt CAACGCGARGAACCTTACC | agt ACAACACGAGCTGACGAC |

B) Genotypes of mouse IDs.

| Mouse ID | Genotype |
|---|---|
| S650 | Kit +/+ |
| S672 | Kit +/+ |
| S656 | Kit +/+ |
| S666 | Kit +/+ |
| S667 | Kit +/+ |
| S658 | Kit +/+ |
| S654 | Kit W/Wv |
| S664 | Kit W/Wv |
| S642 | Kit W/Wv |
| S670 | Kit W/Wv |
| S655 | Kit W/Wv |
| S665 | Kit W/Wv |
| S18367 | Cpa3 +/+ |
| S18368 | Cpa3 +/+ |
| S18189 | Cpa3 +/+ |
| S18166 | Cpa3 +/+ |
| S18163 | Cpa3 +/+ |
| S18366 | Cpa3 Cre/+ |
| S18190 | Cpa3 Cre/+ |
| S18164 | Cpa3 Cre/+ |
| S18365 | Cpa3 Cre/+ |
| S18165 | Cpa3 Cre/+ |
